# Human adenovirus infection induces the overexpression of CLOCK protein in human lymphoblast cells

**DOI:** 10.1101/2022.10.03.509795

**Authors:** Hui Ding, Chin-Fu Chen

**Affiliations:** San Diego, CA, USA; Sinceris consulting, Clemson, SC, USA

**Keywords:** Circadian rhythm, human adenovirus, CLOCK, lymphoblastoid cells

## Abstract

Circadian rhythms are biological processes that regulate metabolism, the immune system, hormones, behaviors, and other various biological processes in organisms. The molecular underpinning of circadian rhythms is a group of genes which regulate each other in transcription-translation feedback loops as an internal molecular clock in cells. Many factors can affect the circadian clock. Viruses, such as hepatitis virus, influenza virus, human immunodeficiency virus were reported to interplay with circadian rhythms in the host or cell level. Up to date, the relationship between viruses and circadian rhythms and its significance in biology, pharmacology and clinic are not entirely understood. We here report that human adenovirus infection could regulate circadian rhythms in cells. We found that human adenovirus infection induced protein overexpression of a core circadian gene *CLOCK* in human lymphoblast cells. The conditioned medium collected from the infected lymphoblast cell culture was able to infect other lymphoblast cell lines and induced CLOCK overexpression in them. In contrast to the previous studies that showed viral infections dampened the circadian oscillation, we found that the human adenovirus infection increased the amplitude of the circadian oscillation in U2OS cells. To our knowledge, this is the first time that adenovirus infection was found to regulate circadian rhythms in cells.

## Introduction

Circadian rhythms are a group of endogenous biological processes with a period of about 24 hours. These processes are conserved across multiple taxa including bacteria, fungi, plant, and animals (1). The circadian oscillations occur within most organs, tissues, and cells in mammals. Circadian rhythms regulate many biochemical, physiological and even behavioral processes, such as hormone secretion, liver metabolism, blood pressure, immune response, mood, and sleep/wake cycle (2, 3). The molecular basis of circadian rhythm consists of a group of circadian genes that regulate each other in transcriptional-translational feedback loops in each cell (4, 5). *BMAL1* and *CLOCK* are two genes in the center of the feedback loops. Their proteins heterodimerize to induce the transcription of other circadian genes, such as *PER1/2/3, CRY1/2, RORα, REV-ERBα*, and many other circadian-controlled genes through binding to regulatory elements (*e*.*g*., E-box) in their promoters (6, 7). The PER and CRY proteins translocate back into the nucleus and repress the actions of the CLOCK/BMAL1 heterodimer as the negative feedback loop (8, 9). RORα positively regulates, while REV-ERBα negatively regulates, the transcription of *BMAL1* through regulatory elements in its promoter (10). Other feedback loops fine-tune these main loops. Light (11), high concentration of serum (12), melatonin (13) and dexamethasone (14) have been reported to be able to adjust or even reset the molecular clock in cells. Viruses were also found interplaying with the host’s circadian rhythms. Many viruses, like Hepatitis B, Hepatitis C, Human Immunodeficiency Virus (HIV) and Simian Immunodeficiency Virus were reported to perturbate circadian gene expression or circadian activities (15–19). On the other side, deficiency or inhibition of circadian genes, like *BMAL1*, promoted viral replication in the host and increased the susceptibility of the host to virus infections (20, 21). The anti-viral immune responses of the host were also under circadian regulation and the circadian time of viral infection affected virus-associated mortality of the host (22, 23).

In this study, we first reported that human adenovirus (hAdV) infection could also affect circadian rhythms. hAdVs are a group of double-stranded DNA viruses with at least 103 known genotypes (24). They are ubiquitous and most commonly cause respiratory diseases and in rare cases serious illness or death. Approximately 5% of human common colds are due to hAdVs infection. Majority of people would have been infected with one or more serotypes of hAdVs during their early lifetime and built up a lifelong immunity (25). Adenovirus vectors have been widely used as a gene delivery tool to transfer genes into target cells, especially in gene therapy for cancer (26). We found that hAdV infection could significantly elevate the CLOCK expression in lymphoblast cells and altered the circadian oscillation in U2OS cells.

## Materials and methods

### Lymphoblast cells culture and conditioned media

Lymphoblast cell lines were obtained by immortalizing lymphocytes from human peripheral blood samples using Epstein-Barr virus. Lymphoblast cell lines were maintained with RPMI-1640 medium (Corning, Corning, NY, USA), supplemented with 12.5% fetal bovine serum (Atlanta Biological, Lawrenceville, GA, USA), 2 mM L-glutamine, 1% penicillin/streptomycin and 1% MEM non-essential amino acid solution (all three from Sigma-Aldrich, St. Louis, MO, USA) in 37°C and 5% CO_2_. To eliminate the possible time effect of feeding and manipulation on circadian genes, all the cell culture was performed at a fixed time of one day (from 12 PM to 1 PM). All the treatment and cell harvest were also performed at a fixed time in each batch.

The conditioned media were collected from human adenovirus C serotype 5 (hAdV5) infected lymphoblast cell culture or uninfected lymphoblast cell culture (as control) 3 days after changing media. Then they were filtered with 0.2 μm PVDF filters and stored at -20°C for future use.

### Conditioned media infection

Four uninfected lymphoblast cell lines were used in the conditioned media infection. Total 4 × 10^6^ cells of each line were pelleted and resuspended with either 4 mL of hAdV5 conditioned medium or control conditioned medium into a T25 tissue culture flask, and then incubated in an incubator with 37°C and 5% CO_2_. Four mL of fresh medium was added to each flask on the third day and all the cells were harvested on the fifth day after the infection. Half of the cells were used for DNA and total RNA isolation. The other half were used for whole cell lysate preparation.

### SDS-PAGE, protein visualization and Western blotting

Lymphoblast cells were lysed using RIPA buffer (150 mM NaCl, 1% NP-40, 0.5% sodium deoxycholate, 0.1% SDS, 50 mM Tris pH8.0) supplemented with protease inhibitors. The whole cell lysates were mixed with Laemmli sample buffer (BIO-RAD, 1610737) and SDS, and then boiled for 10 minutes. Precast 10% polyacrylamide gels (BIO-RAD, 4561033) were used in SDS-PAGE. Coomassie blue was used to visualize the proteins in the protein gel. A precast 10% polyacrylamide stain-free gel (BIO-RAD, 4568035) was also used to visualize proteins in the gel. This gel contains trihalo compounds which react with tryptophan residues in the proteins and produce fluorescence in the presence of UV-light. A BIO-RAD ChemiDoc MP Imaging System was used in the protein visualization.

Anti-CLOCK (sc-25361), anti-BMAL1 (sc-365645) and anti-GAPDH (sc-32233) used in the Western blotting were purchased from Santa Cruz Biotechnology (Santa Cruz, CA, USA). The bands on the film were quantified using ImageJ (NIH).

### Protein mass spectrometry

After staining with Coomassie blue, the strong band in the gel was cut and sent to the Proteomics & Mass Spectrometry Facility at University of Massachusetts Medical School for protein mass spectrometry analysis. All the identified peptides were matched to all the protein entries in Swiss-Prot database to identify the proteins. Then the identified proteins in the sample were ranked based on the normalized quantitative value.

### PCR and RT-qPCR

The DNA of lymphoblast cell lines was isolated by using a DNA precipitation method. Briefly, cells were collected, lysed using 90% lysis buffer (75 mM NaCl, 25 mM EDTA) and 10% sarkosyl-proteinase K solution (2 mg proteinase K/mL sarkosyl solution), and rotated overnight at 4°C. Then saturated sodium chloride solution and ice-cold isopropanol were added and mixed. When DNA precipitated, DNA was spooled with a sterile Pasteur glass pipet and transferred to TE buffer (10 mM Tris-HCl, 1 mM EDTA, pH 8.0) to dissolve overnight. Primers used for PCR are listed in Supplementary Table S1.

Total RNA was isolated using the RNeasy Mini RNA isolation kit (Qiagen, Valencia, CA, USA) according to the manufacturer’s instructions. Reverse transcription was performed using Verso cDNA kit (Thermo Fisher Scientific, Waltham, MA, USA). Real-time quantitative PCR (RT-qPCR) for quantification of mRNA was conducted using iQ SYBR Green Supermix and a BioRad CFX96 Real-time PCR System (BioRad, Hercules, CA). The qPCR primers were designed using the Primer3 online tool, and the sequences was listed in Supplementary Table S1. Our preliminary analyses suggested that the level of mRNA of the glyderaldehyde-3-phosphate dehydrogenase (*GAPDH*) gene was relatively similar across most of lymphoblast cell lines tested. Thus, it was used as the internal reference gene. Each sample was analysed in three replicates.

### Luciferase assay and conditioned medium treatment

A U2OS cell line (human bone osteosarcoma epithelial cells) with a luciferase reporter was provided as a gift from Dr. Luciano DiTacchio (Medical Center, University of Kansas). The luciferase reporter was driven by the *mBmal1* promoter (27). U2OS cells were maintained in DMEM (Invitrogen, Carlsbad, CA, USA) plus 10% FBS, 1% Penicillin/Streptomycin and 2 μg/mL puromycin. A total of 4 × 10^5^ U2OS cells were seeded in each well of a 96-well plate (white, opaque) and incubated in 37°C and 5% CO_2_ overnight. A final concentration 100 nM of dexamethasone was added to synchronize the cells for two hours. After synchronization, the medium in each well was replaced with Phenol Red-free DMEM, 10% FBS, D-Luciferin (final concentration 100 μM) and the proportional (10%, 20%, 30%, and 40%) conditioned medium from control (uninfected) lymphoblast cells or hAdV5 infected lymphoblast cells. Each condition had five replicate wells. The final volume was 200 μL in each well. After a two-hour incubation in 37°C and 5% CO_2_, the plate was sealed with a transparent membrane and subjected to real-time bioluminescence measurement (Spark multimode microplate reader, Tecan, Switzerland) in every 15 minutes for 84 hours in 37°C.

### Cell viability

A total of 4 × 10^5^ U2OS cells were seeded in each well of a 96-well clear plate in 37°C and 5% CO_2_ overnight, and then processed the same conditions of synchronization, conditioned medium treatment and membrane sealing as those in the luciferase assay. After 26 hours, the supernatant in each well was collected for suspension cell counting. The adherent cells on the bottom of the plate were detached using Trypsin-EDTA and collected for adherent cell counting. An automated cell counter (TC10, Bio-Rad) and trypan blue were used for cell counting and viability measurement.

### Data processing and statistical analysis

We used a standard curve method to calculate the relative copy numbers of the genes of interest and reference gene (*GAPDH*) in the RT-PCR experiments. The Student’s *t*-test or one-way analysis of variance (ANOVA) and then Tukey’s HSD test were used to evaluate the significant difference.

## Results

### CLOCK overexpression and human adenovirus infection were found in two lymphoblast cell lines

In our study on the expression of circadian genes in patients with autism spectrum disorder (ASD), we unexpectedly found that the lymphoblast cell lines from two brothers with ASD displayed high expression of CLOCK protein, but not its molecular partner BMAL1(Fig 1A). When running the lysates of the two lymphoblast cell lines on SDS-PAGE, a dark band at about 120 kDa in each of them drew our attention (Fig 1B). The dark band was cut and analysed by protein mass spectrometry. The main protein in the band was found to be hexon, a major coat protein in human adenovirus C serotype 5 (hAdV5) (protein identification probability: 100%, percentage sequence coverage: 58.2%). We then used PCR to detect two genes of hAdV5 *Hexon* and *E1A* in the DNA of the two lymphoblast cell lines and both were positive (Fig 1C). The PCR amplicon of *Hexon* was sequenced and it matched 97% of the sequence of *Hexon* gene in hAdV5 (NCBI Reference Sequence: AC_000008.1) (data not shown). We hence concluded that the two lymphoblast cell lines were infected by hAdV5.

**Fig 1.**
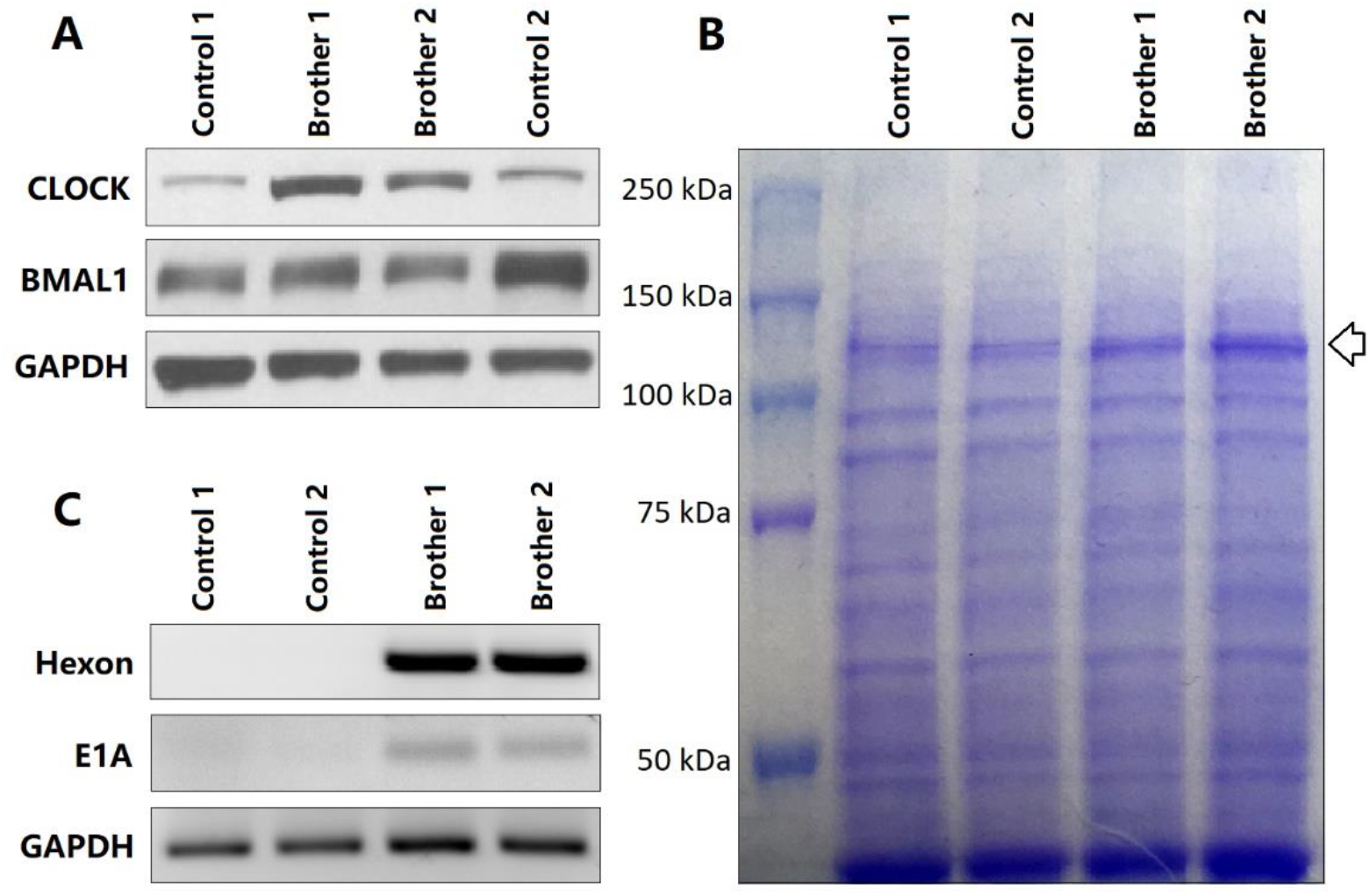
CLOCK overexpression and human adenovirus infection were found in two brothers’ lymphoblast cell lines. Two control lymphoblast cell lines and the two brothers’ lymphoblast cell lines were cultured and prepared cell lysates for (A) Western blotting analysis of CLOCK and BMAL1, (B) SDS-PAGE Coomassie staining, and isolated DNA for (C) PCR analysis of hAdV5 genes *Hexon* and *E1A*. In (B), the arrow indicates the position of the dark bands.

We tested an additional 40 lymphoblast cell lines from normal controls and 40 lymphoblast cell lines from ASD patients in our lab and did not detect the DNA of the hAdV5 genes *Hexon* and *E1A*, nor did we find the significant CLOCK overexpression in them (data not shown).

### The hAdV5 conditioned medium was infectious for lymphoblast cells

To investigate whether the infected lymphoblast cells release hAdV5 viruses into the culture medium so that the conditioned medium is infectious for other lymphoblast cells, we performed the following experiments. Conditioned medium from one of the two hAdV5-infected lymphoblast cell lines was collected and used to treat four uninfected human lymphoblast cell lines from normal controls. Conditioned medium from an uninfected human lymphoblast cell line from a normal control was collected to treat the same four cell lines as controls. After a 5-day treatment, dark bands appeared at around 120 kDa, the size of hexon protein, on the protein gel in all the samples of the lymphoblast cell lines treated with hAdV5 conditioned medium (Fig 2A). The DNA of all the lymphoblast cell lines were also isolated. The two hAdV5 genes, *Hexon* and *E1A*, were identified in the DNA of the lymphoblast cell lines treated with hAdV5 conditioned medium, but not in the controls (Fig 2B). The observations suggested that the conditioned medium collected from the hAdV5 infected lymphoblast cells was infectious for other lymphoblast cell lines.

**Fig 2.**
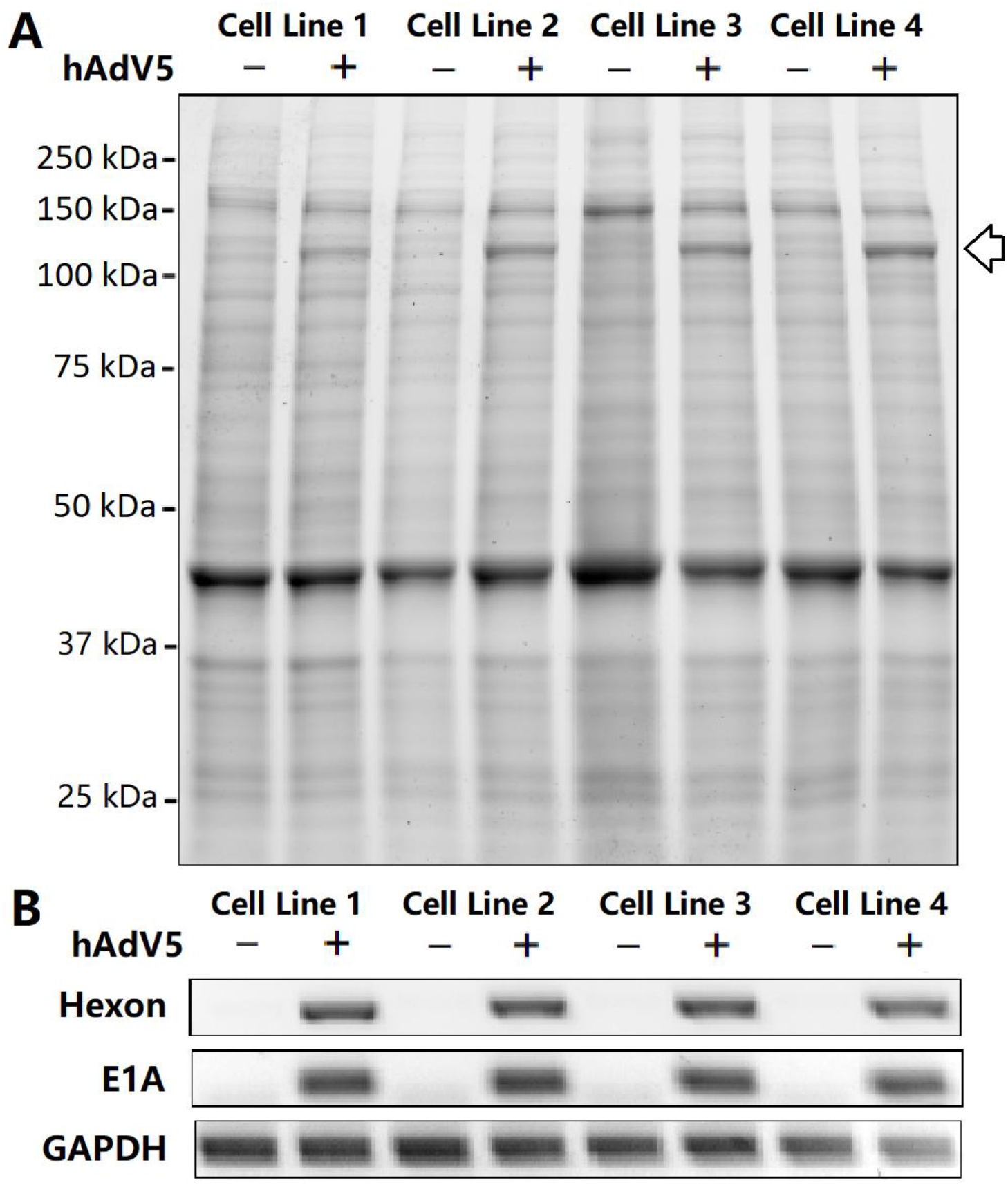
hAdV5 conditioned medium infected other lymphoblast cell lines. Four uninfected lymphoblast cell lines were treated with conditioned medium either from uninfected or hAdV5 infected lymphoblast cell lines for five days. (A) Trihalo SDS-PAGE analysis of the cell lysates from these treated cell lines. The arrow indicates the position of the dark bands related to the hAdV5 hexon protein. (B) PCR analysis of hAdV5 genes *Hexon* and *E1A* in the DNA of the treated cell lines.

### The hAdV5 infection promoted the CLOCK protein expression

To address the question whether the conditioned medium infection induced CLOCK overexpression in the lymphoblast cell lines, we performed Western blotting analysis of CLOCK and BMAL1 in these lymphoblast cell lines treated with control or hAdV5 conditioned media. The level of CLOCK protein increased more than 11-fold in the lymphoblast cell lines treated with hAdV5 conditioned medium compared to the ones treated with control conditioned medium (*p* value = 0.02). BMAL1 also increased about 47% in the lymphoblast cell lines treated with hAdV5 conditioned medium, although it was not statistically significant (*p* value = 0.18) (Figs 3A and B). The above observations suggested that hAdV5 infection remarkably promoted the expression of CLOCK protein in lymphoblast cells.

**Fig 3.**
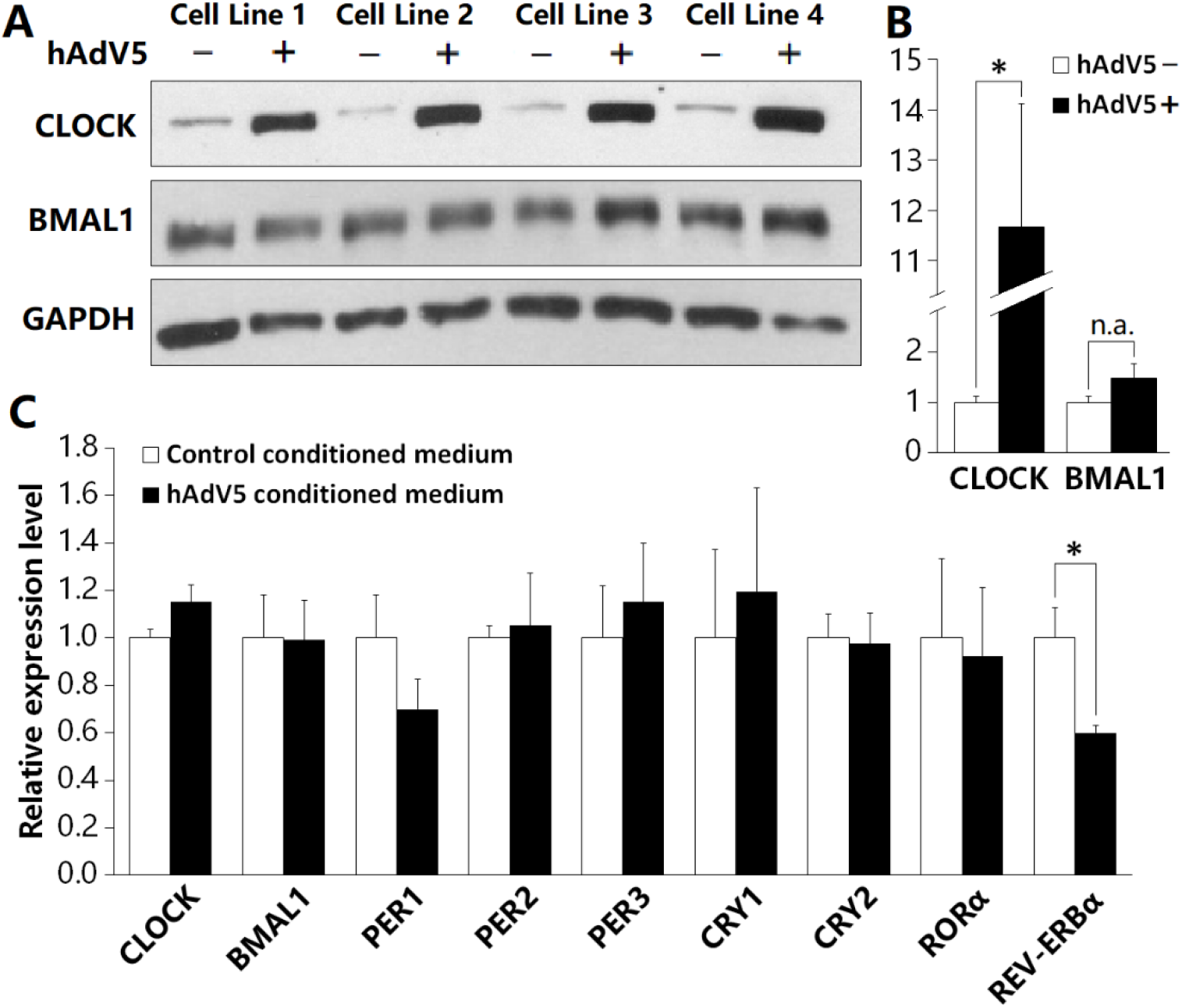
Effect of hAdV5 conditioned medium treatment on circadian genes expression in lymphoblast cells. Four uninfected lymphoblast cell lines were treated with conditioned medium from uninfected or hAdV5 infected lymphoblast cell lines for five days. (A) The whole cell lysates of all the lines were prepared for a Western blotting analysis of CLOCK and BMAL1. GAPDH was used as reference protein. (B) The intensities of the CLOCK and BMAL1 bands in (A) were quantified and normalized to that of GAPDH bands. (C) The total RNA of all the cell lines were isolated for analysis of circadian genes expression using RT-qPCR. GAPDH was used as internal reference gene. The values are mean ± SEM (n = 4). Asterisks indicate *p* ≤ 0.05.

### The hAdV5 infection affected the transcription of circadian genes

To investigate if the hAdV5 infection affected the transcription of the circadian genes, the mRNA levels of nine core circadian genes were measured using RT-qPCR in these lymphoblast cell lines treated with control or hAdV5 conditioned media. We found the mRNA level of *CLOCK* increased modestly by 15% in the lymphoblast cells treated with hAdV5 conditioned medium compared to the controls, but no difference in the mRNA level of *BMAL1* in the lymphoblast cells treated with hAdV5 or control conditioned medium (Fig 3C). Among the negative regulators of *CLOCK* and *BMAL1*: *PER1, PER2, PER3, CRY1, CRY2*, and *REV-ERVα*, only *PER1* and *REV-ERVα* were noticeably decreased by 30% (*p* value = 0.066) and 40% (*p* value = 0.050) in the lymphoblast cells treated with hAdV5 conditioned medium. *PER3* and *CRY1* even increased slightly in the lymphoblast cells treated with hAdV5 conditioned medium. There was no significant change in the mRNA level of *RORα*, the positive regulator of *CLOCK* and *BMAL1*, in the hAdV5 conditioned medium treated lymphoblast cells (Fig 3C).

### The hAdV5 infection altered circadian oscillation in U2OS cells

To use a luciferase reporter driven by a circadian gene promotor can transform the circadian oscillation in cells into a bioluminescent signal. We used a U2OS cell line containing a luciferase reporter driven the *mBmal1* promoter (27) and different proportions of conditioned medium from the hAdV5 infected lymphoblast cells or uninfected cells was added to the culture of U2OS cells. The bioluminescence which reflects the circadian oscillation in U2OS cells was monitored and recorded. In the first circadian period, the amplitudes of the circadian oscillation in the U2OS cells treated with the hAdV5 conditioned medium were 16.4% higher than those in the cells treated with control conditioned medium, and 13.7% higher than those in the cells without any conditioned medium treatment. The oscillation amplitudes in cells with the control conditioned medium and without conditioned medium treatment were not statistically different (Figs 4A and B). The time to the first peak was also significantly extended in the U2OS cells treated with hAdV5 conditioned medium. The cells treated with hAdV5 conditioned medium needed an additional 2.47 hours to get the first peak compared to the cells without conditioned medium treatment. The time to get to the first peak in the cells with control conditioned medium also increased by 1.74 hours compared to that in the cells without conditioned medium treatment (Figs 4A and B).

**Fig 4.**
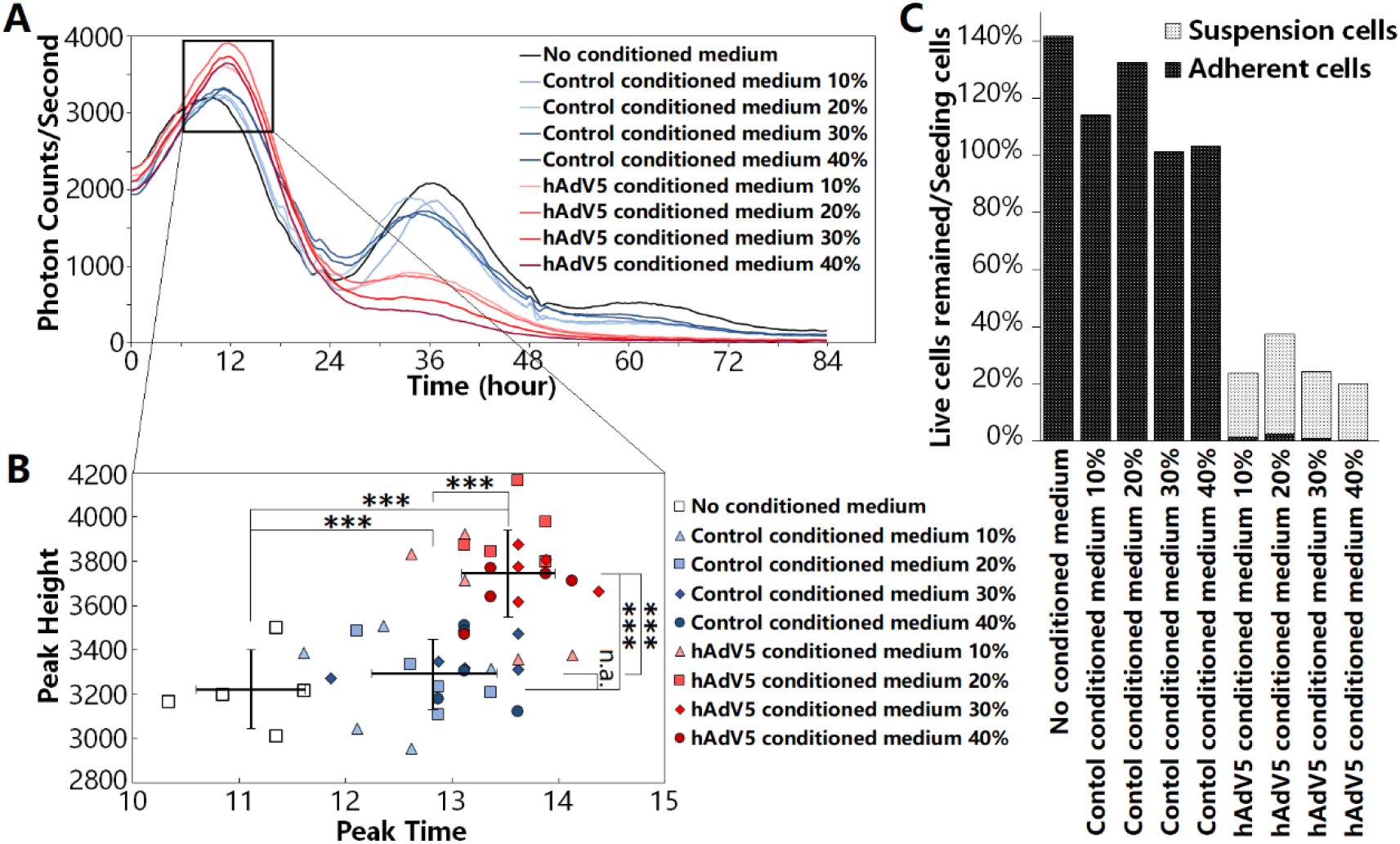
Effect of conditioned medium treatment on circadian oscillation and viability of U2OS cells. (A) Conditioned medium from uninfected or hAdV5 infected lymphoblast cell lines were used to treat U2OS cells with a luciferase reporter driven by mBmal1 promoter in 10%, 20%, 30% and 40% of the final volume. Each treatment had five replicates. Two hours after the conditioned medium added, the luminescent signals were started to be monitored. Each line standards for the average of the five replicates. (B) The time and height (luminescent signal strength) of the first peak in all the replicates and conditions in (A) were extracted and plotted. The bars and crossing points indicate standard deviations and means of no conditioned medium treatment group, control conditioned medium treatment group and hAdV5 conditioned medium treatment group. (C) The viability of U2OS cells was examined 26 hours after the conditioned medium added (n = 5). In each of the bars, the light portion indicates the viability of the suspension cells (which is close to zero in the no conditioned medium treatment and control conditioned medium treatments), and the dark portion indicates the viability of the adherent cells (which is very small in the hAdV5 conditioned medium treatments). The three asterisks indicate significance, *p* value < 0.001.

In the second circadian period, the cells treated with the control conditioned medium and the cells without any conditioned medium treatment kept oscillating. However, the cells with hAdV5 conditioned medium treatment failed to generate the second crest. When inspected under the microscope, the cells treated with hAdV5 conditioned medium were found detached from the culture plate. To investigate if the U2OS cells were still alive, the viability of the adherent and suspension cells was determined respectively after a 26-hour incubation in each condition. The results showed that only 0.5% to 2.7% of the cells were still alive and adherent on the culture plate in the treatment with hAdV5 conditioned medium in different proportions, and totally 20.0% to 37.5% of cells were still alive (Fig 4C). In contrast, the cells treated with control conditioned medium maintained the seeding number of cells or grew more, and there were nearly no cells found in the supernatant. The cells without treatment of any conditioned medium increased the cell number by 41.7% (Fig 4C). These observations suggested that conditioned medium collected from hAdV5 infected lymphoblast cells were detrimental to U2OS cells.

## Discussion

We do not know how the two lymphoblast cell lines were infected by hAdV5. All the lymphoblast cell lines used in this report were established by immortalizing lymphocytes from human peripheral blood with Epstein-Barr virus following standard procedures. The two hAdV5 infected lymphoblast cell lines were created at different times from the blood samples of two brothers with autism spectrum disorder (ASD). We conjecture that the two brothers might have already been infected by hAdV5 at the time of collecting their blood samples. However, we don’t believe that the consequent CLOCK overexpression and altered circadian rhythm are related to the pathogenesis of ASD. The lymphoblast cell lines from non-ASD controls could also be infected by the hAdV5 to overexpress CLOCK. In our lab, we also frequently use HEK293 cells purchased from ATCC (the American Type Culture Collection). HEK293 cells were generated by transfecting human embryonic kidney cells with sheared hAdV5 DNA in1973, but there is no evidence suggesting that HEK293 cells can produce hAdV5 viruses and infect other cells (28). Furthermore, we used the conditioned medium from HEK293 cells to treat lymphoblast cell lines and no infectivity of adenovirus was observed (data not shown).

The overexpression of CLOCK protein was one of the main effects we found in the hAdV5 infection. More than 11-fold increase of CLOCK was found in lymphoblast cells after hAdV5 infection. There were also some other modest changes, such as increase in *CLOCK* mRNA and BMAL1 protein, and decrease in *PER1* and *REV-ERVα* mRNA. The other circadian genes in the circadian machinery remained relatively intact. Most of the circadian genes are transcription factors. About 5% to 25% of mammalian genes were identified circadian oscillating in transcription, and most of them are associated with the areas of metabolism and immunity (29). Viruses depend on the machinery and resources of the host cell to synthesize their own DNA, RNA and proteins to reproduce themselves, and fight with the host’s immune system. The circadian genes and system might be the good targets for viruses to exploit and manipulate to benefit their replication and survival.

Besides being a transcription regulator in circadian rhythms, CLOCK protein is also a histone acetyltransferase (HAT), and BMAL1 enhances its HAT function (30). The CLOCK/BMAL1 dimer with the HAT activity can modulate the chromatin structure and make it much easier to access and activate the target genes. HAT also plays critical roles in the gene transcription of dsDNA viruses. When viral dsDNA enters the nucleus of the host cell, histones trend to bind and attempt to silence it. CLOCK, as an HAT, can disrupt this suppression and facilitate the expression of the virus genes. For example, CLOCK was found essential in the viral transcription of herpes simplex virus 1 (HSV-1). HSV-1 also helped to stabilize CLOCK and block its degradation in the infected host cells (31, 32). hAdV5 is a dsDNA virus and may also elevate and stabilize CLOCK to help its viral gene expression.

We also found that the circadian oscillation was altered after hAdV5 infection in U2OS cells. The hAdV5 infection significantly increased the amplitude of the first circadian period. This may be related to the overexpression of CLOCK protein. Some other studies also found that virus could perturb circadian rhythm, but most of the perturbance were reducing the oscillation amplitudes. For instance, virus infection or chronic expression of viral protein would cause reduced amplitude of the circadian oscillation in blood pressure and heart rate of human (33), in temperature and locomotor activity of monkey (15), and in locomotor activity of mouse (34, 35). Only in one study, Dengue virus infection increased the amplitude of mosquito’s rhythmic locomotor activity, which facilitated the dissemination of the virus (36). The hAdV5 infection time also extended the time to get to the first peak, which could be caused by the lack of nutrients or the effect of some metabolic products in the conditioned medium, since the control conditioned medium treatment also caused the extended peak time.

The sine wave in the first period was intact revealing that the circadian feedback loops were well maintained at the beginning of the hAdV5 infection. However, the U2OS cells were not able to survive the hAdV5 infection for a long time like lymphoblast cells. Most of U2OS cells were suspended or dead after the first period which should be the reason that they failed to oscillate the second circadian crest. Because of cell death, the cell lysate could not be collected to determine the CLOCK protein level in U2OS cells infected by hAdV5. Lymphoblast cells can survive the hAdV5 infection and maintain a high level of CLOCK. It can be speculated that the circadian oscillation in lymphoblast cells may be altered and maintained for a long time by the hAdV5 infection. Due to the difficulties to deliver genes into lymphoblast cells, we cannot investigate the circadian rhythm changes in the hAdV5 infected lymphoblast cells.

Adenovirus has been widely used as a gene delivery vehicle in scientific studies and gene therapy clinic trials (26, 37), including in some circadian rhythm studies (38). Although the adenovirus used for gene delivery is usually replication-incompetent, its perturbation to circadian rhythm should still be regarded with caution based on our study. The study for the interplay between viruses and the host’s circadian rhythms is currently at its infancy. Our report about the effect of hAdV5 infection on circadian rhythms may lead to better understanding of their relations and some guiding significance in study and clinical applications.

## Acknowledgements

The experimental work was primarily done while the authors were at the J.C. Self Research Institute of Human Genetics, Greenwood Genetic Center, Greenwood, South Carolina. We are grateful for the support provided by the Greenwood Genetic Center. We also thank Dr. Charles E. Schwartz at Greenwood Genetic Center and Dr. Yanzhang Wei at Department of Biological Sciences, Clemson University for reviewing a previous version of the manuscript. We thank Dr. Luciano DiTacchio (Medical Center, University of Kansas) for the U2OS cells with the luciferase reporter.

## Notes

### Competing Interest Statement

The authors have declared no competing interest.

